# Land use influences human mental and respiratory health in a conservation priority area in southeastern Brazil

**DOI:** 10.1101/2022.09.22.508330

**Authors:** Matteus Carvalho Ferreira, Rodrigo Lima Massara, Marcelo de Ávila Chaves, Bruno Eduardo Fernandes Mota, Flávio Henrique Guimarães Rodrigues

## Abstract

Human activities generate negative environmental impacts that can compromise ecological processes and ecosystem services and thus, influence human health. We assessed how natural and altered areas affect human respiratory and mental health in one of the largest mining provinces in the world, the Quadrilátero Ferrífero (QF), in southeastern Brazil. We used a model selection approach to evaluate socioeconomic and environmental factors that would influence hospitalization rates for mental and behavioral disorders, as well as for respiratory diseases in 22 municipalities in QF. Municipalities with higher rates of urbanization had higher rates of hospitalizations for mental and behavioral disorders. Conversely, the adult population (15-59 years old) of both gender and the elderly female population (60 years old or more) presented lower rates of hospitalization for respiratory diseases in municipalities with a higher rate of urbanization, suggesting a greater ease of access to prophylactic measures of the population located in these municipalities compared to those with a lower rate of urbanization. Municipalities with larger urban forest areas had lower rates of hospitalization for respiratory diseases in the young (0-14 years) female population, while municipalities with larger mining areas had higher rates. The elderly male population (60 years or older) also had higher rates of hospitalization for respiratory diseases in municipalities with larger mining areas. Our findings show important ecosystem services provided by urban forests and highlight impacts on health, in different segments of the population, due to anthropogenic changes in the landscape.

**Highlights:**

- We present a multiscale method to determine factors that influence human health.
- Demographic groups are differently influenced by socio-environmental variables.
- Urbanization rate is associated with worse mental health of human population.
- Forest in urban areas is associated with better respiratory health for children.
- Mining is associated with worse respiratory health for children and elderly.

**Graphical Abstract:** 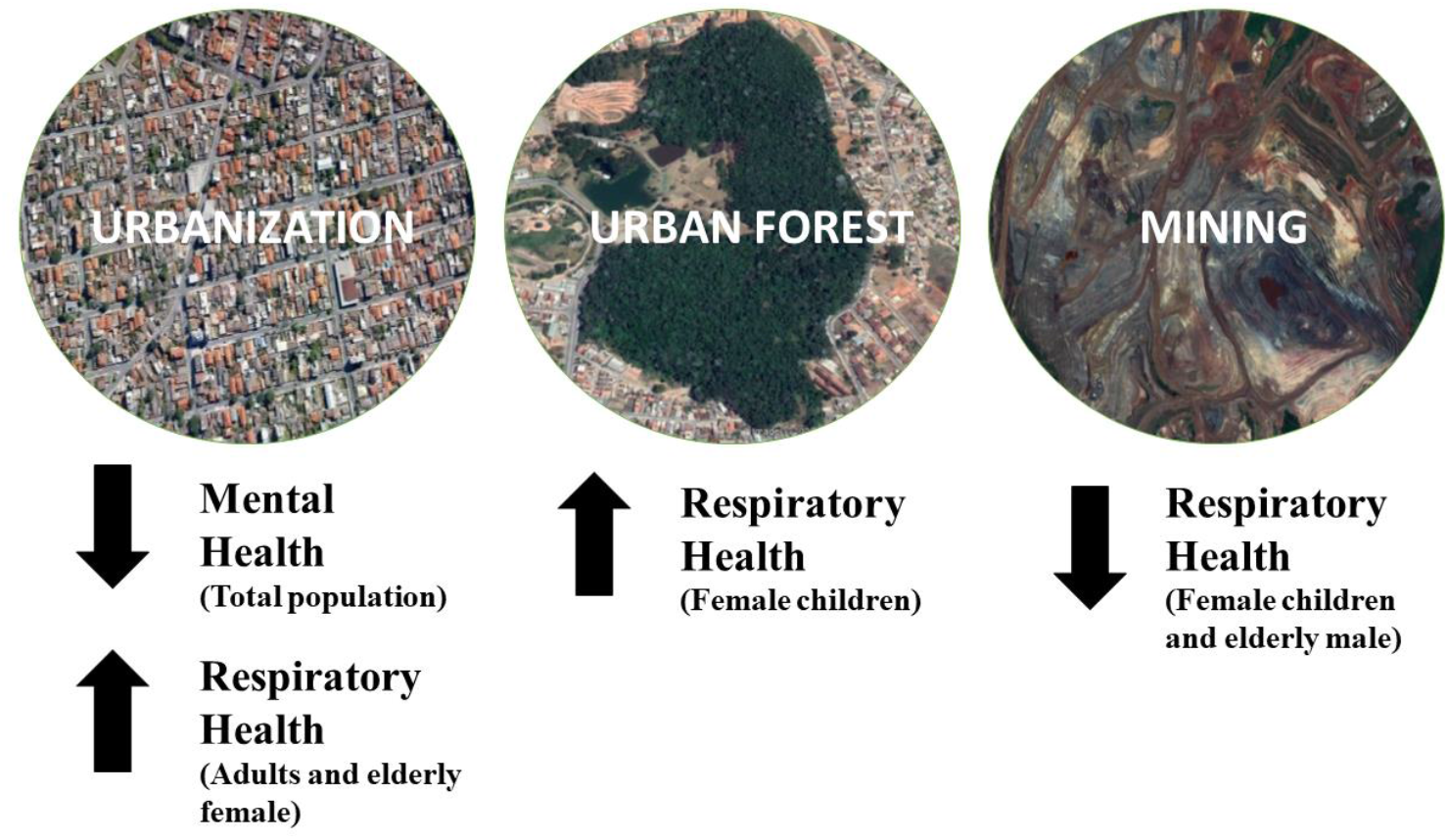

## Introduction

Natural environments provide different ecosystem services to society, ranging from food, water, and air quality, to the feeling of well-being and disease reduction (Díaz et al., 2018). Despite this services, natural systems are being degraded to an unprecedented extent in human history, resulting in a climate emergency scenario (Ripple et al., 2019). Environmental impacts, caused by climate change and human land use and occupation, end up compromising planetary health and threatening human health in ways that are still uncertain (Whitmee et al., 2015). Therefore, it is becoming increasingly urgent to adopt an integrated and multidisciplinary vision involving human health and ecosystem health from a perspective of the “One Health” paradigm (Ryu et al., 2017).

The relationship between human health and the environment has been highlighted in several recent global studies and reports (Bratman et al., 2019; Frumkin et al., 2017; Romanelli et al., 2015; Whitmee et al., 2015). These studies show the positive effects of natural areas on human health, as well as warn about the negative impacts of anthropic activities, which need to be internalized to define efficient public health policies. It is necessary to understand the socio-environmental factors that relate to human health, to account for the depreciation of natural capital and its socioeconomic costs, integrating environmental health into the budget for the prevention of diseases and the promotion of equal and universal collective health (Whitmee et al., 2015).

Encouraging contact with nature is a form of prevention and treatment of several groups of diseases with excellent cost-benefit ratio, presenting low financial costs when compared to conventional medical interventions (Frumkin et al., 2017). At the regional and local level, the trend towards increased urbanization and reduced contact with natural environments may pose a risk to human health (Cox et al., 2018; Mills et al., 2019; Whitmee et al., 2015), especially for mental health (Bratman et al., 2019).

Respiratory diseases are among the main causes of hospitalization in the world (Romanelli et al., 2015), while mental and behavioral disorders are one of the fastest-growing diseases on planet and are those that have the highest associated economic impact when considered direct and indirect costs (Trautmann et al., 2016). In Brazil, in 2010, there were 1,452,505 hospitalizations for respiratory diseases at a cost of R$ 1,192,711,452.26 (or R$ 1,989,700,208.77 in adjusted values for 2020 - R$1 = US$0.19, August 3^rd^, 2020) and 159,746 hospitalizations for mental and behavioral disorders at a cost of R$ 172,046,034.84 (or R$ 287,009,930.85 in adjusted values for 2020), this considering only the direct costs for the Unified Health System (acronym in Portuguese, SUS; SUS, 2019). Understanding the factors that influence these rates is necessary, not only from a humanitarian point of view but also from an economic and territorial planning perspective (Pattanayak et al., 2017).

We hypothesized that different socioeconomic and environmental factors would influence hospitalization rates for respiratory diseases and for mental and behavioral disorders (hereafter, mental illness). We also expected that these variables would influence hospitalizations differently according to the age group and gender. This separation was conducted to identify the factors that could influence hospitalization rates differently since studies show that vulnerabilities to diseases can be influenced by gender (Buvini et al., 2006; Sakiani et al., 2013) and by age (Akha, 2018; Nikolich-Žugich, 2018). Specifically, we expected a positive correlation of urban and mining areas in the landscape with rates of human hospitalization for respiratory and mental illness, while forest areas would reduce these hospitalization rates. As other factors can affect hospitalization rates, we also assessed whether human vaccination coverage and average rainfall (i.e., precipitation) would influence the rates of human hospitalization for respiratory diseases. Likewise, we assessed whether medical availability, urbanization rates and the human development index of each municipality would influence hospitalization rates for respiratory and mental illness. The study area is one of the largest mining provinces in the world, the Quadrilátero Ferrífero, located in Southeastern Brazil.

## Materials and methods

### Study area

The study region was the Quadrilátero Ferrífero (QF), an area of approximately 6,500 km² located in the south-central portion of the state of Minas Gerais (MG), southeastern Brazil (Figure 1). The region, which is part of the Serra do Espinhaço Biosphere Reserve and Mata Atlântica Biosphere Reserve (UNESCO), has a great heterogeneity in the land uses, with emphasis on mining and urbanization. QF is also considered one of the most important regions for the Brazilian biodiversity, with the dominance of the Cerrado and the Atlantic Forest domains (i.e., biodiversity hotspots), in addition to the Campos Rupestres ecosystems (Silveira et al., 2015). Despite centuries of human occupation and mining activities (i.e., 55% of the region is composed of agribusiness, mining, and urban areas), the region is still home to great biodiversity, with about 45% of the territory composed of native vegetation (Sonter et al., 2014). Despite this, the increase of mining activities and urbanization threatens the ecosystem services in the QF, especially those related to the quality of habitats, carbon stocks and sediment retention and thus, compromising water availability, the combat against climate change, and biodiversity conservation (Duarte et al., 2016).

**Figure 1:**
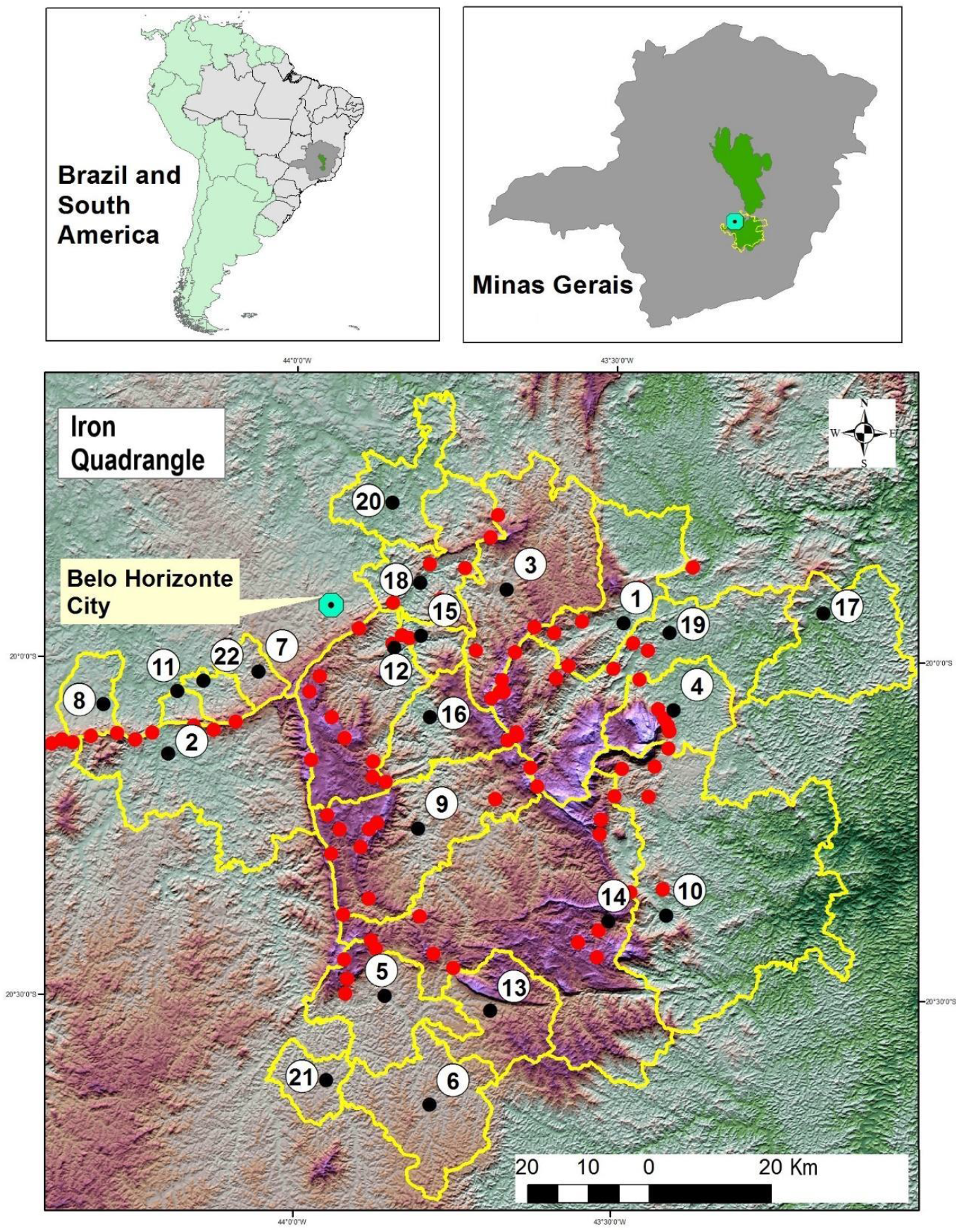
Map of location and details of the study area. The yellow lines indicate the limits of the 22 municipalities selected and evaluated and the red dots show the main active mining operations. The original limits of the Serra do Espinhaço Biosphere Reserve are highlighted in green in the insert maps of Brazil and Minas Gerais. The municipalities of the Quadrilátero Ferrífero considered were: Barão de Cocais (1), Brumadinho (2), Caeté (3), Catas Altas (4), Congonhas (5), Conselheiro Lafaiete (6), Ibirité (7), Igarapé (8), Itabirito (9), Mariana (10), Mário Campos (11), Nova Lima (12), Ouro Branco (13), Ouro Preto 14), Raposos (15), Rio Acima (16), Piracicaba River (17), Sabará (18), Santa Bárbara (19), Santa Luzia (20), São Brás do Suaçuí (21) and Sarzedo (22).

### Data collection

We compiled information from 22 municipalities (i.e., sample unit) of the QF (Figure 1), with a total population in 2010 of 1,191,249 people. Those municipalities with the highest urbanization rates (urban population / total population) were selected, considering the threshold of ≥ 75% (IBGE, 2010). This threshold was adopted to allow analysis at different scales of landscape within the same municipality: the urban centers, the region surrounding these centers, and the area of the city itself (see Data analysis for more details). All data collected were for the year 2010 because of the greater availability of socioeconomic and demographic data due to the Census of the Brazilian Institute of Geography and Statistics (IBGE, 2010).

To assess the population health of each municipality, we collected data on hospitalization rates, average hospitalization time, and total hospitalization costs for respiratory diseases and mental illness using the Brazilian DATASUS platform (SUS, 2019a) of the Brazilian Unified Health System (SUS). All respiratory diseases (J00-J99) included in Chapter X of the International Statistical Classification of Diseases and Related Health Problems - Tenth Revision (WHO, 2004) were considered. For mental illness (Chapter V; WHO, 2004) were considered the following: mental and behavioral disorders due to the use of alcohol and other psychoactive substances (F10-F19), mood disorders (affective) (F30-F39), neurotic and stress-related and somatotropic disorders (F40-F48). The number of hospitalizations in one municipality refers to every individual residing in that municipality who was hospitalized, regardless whether or not he / she was admitted to a Health Unit in another municipality.

### Data analysis

Due to the low number of records for mental illness, we only analyzed the hospitalization rate considering the total population. For respiratory diseases, we analyzed the hospitalization rate in six demographic groups as follows young (0-14 years old) female and male, adult (15-59 years old) female and male and elderly (≥ 60 years old) female and male. The young age group (0-14 years) was defined based on the lower limit of the economically active population (< 15 years), while the elderly age group (≥ 60 years) was defined based on the Brazilian Elderly Statute (Federal Law 10.741/2003). The hospitalization rate was then divided by the population of each demographic group of each municipality to have the number of hospitalizations *per capita*.

To assess whether medical availability would influence hospitalization rates, we obtained information on the number of doctors from each municipality in the SUS Health Information Notebooks (SUS, 2019). For the analysis of respiratory diseases, we used the number of general practitioners, family doctors, and pediatricians, being the latter considered only for the age group for up to 14 years. For the analysis of mental illness, we used the number of general practitioners, family doctors, psychiatrists, psychologists, and pediatricians, being the latter considered only for the age group up to 14 years. The number of doctors was divided by the population of each demographic group of each municipality to obtain the number of doctors *per capita*. Due to the limited availability of data, it was not possible to obtain the number of doctors for the rural and urban population separately.

To verify whether or not the urbanization rate of the municipalities would influence both rates of hospitalization, we obtained data on the number of inhabitants for each demographic group, as well as the respective urbanization rates according to the 2010 IBGE census (IBGE, 2019). To assess whether or not factors related to *per capita* income, longevity, or education in the municipality would affect both hospitalization rates, we obtained the Municipal Human Development Index (MHDI) values in the Human Development Atlas of Brazil (Atlas Brasil, 2019).

To assess how the landscape composition, especially native tree vegetation and mining enterprise would influence both hospitalization rates, we performed a multi-scale spatial analysis in ArcGIS v.10.3 (Esri, 2011). Specifically, we evaluated the composition of the landscape in each of the 22 municipalities considering: 1) the area of the municipality (FJP, 2018); 2) the area of a 3 km radius buffer from the perimeter of the urban centers (i.e., region bounded by a set of buildings visible in satellite images) that encompass the municipality; and 3) the area of the urban centers within the municipality. The 3 km radius buffer was chosen based on a study that evaluated the influence of landscape diversity on the respiratory health of the Australian population (Liddicoat et al., 2018). The perimeter of the urban centers of each municipality was manually delimited from LANDSAT 5 satellite images for the year 2010 (INPE, 2011) and from Google Earth Pro satellite images (Google, 2019) retroactive to the same period. We also used the areas delimited by the perimeter of urban centers to assess if the size of urban centers in cities would influence both hospitalization rates.

The natural forest areas (i.e., Cerrado and Atlantic Forest domain) were extracted from land use and occupation maps from the 3.1 collections of the MapBiomas Project (MapBiomas, 2019) for the year 2010, with the resolution of 30 meters. The mining areas (ditches, dams, waste piles, and processing plants) were measured manually using LANDSAT 5 satellite images (INPE, 2011) and Google Earth Pro satellite images (Google, 2019) for the same period or the nearest year. As the resolution of LANDSAT 5 images is 30 m, the forest areas inside the cities refer mainly to parks, urban forests, and other regions of tree density.

The forest and mining areas were then evaluated at the three different landscape scales. We used the proportion of forest areas in the municipality to assess if more forested municipalities have lower rates of hospitalization. For the other scales (urban centers and 3 km buffer area surrounding these centers), we divided the area of forests present by the urban population of each demographic group, obtaining the area of forest *per capita*. Mining was only present within the scales of the municipality and the 3 km buffer area. Mining had its area divided by the resident population in each of these scales and for each demographic group, obtaining the mining area *per capita*.

Specifically, for respiratory health, we added two variables in the analysis. To assess whether immunization would influence the hospitalization rate, we obtained from DATASUS for the 1996 and 2010 (SUS, 2019) interval, the vaccination coverage data for the following vaccines: Pneumococcal, Measles, Dual Viral, Triple Viral (MMR), Triple Bacterial (DTP), *Haemophilus influenzae* type B, Tetra, and Penta. Due to the limited availability of data, it was not possible to obtain immunization rates for the rural and urban populations separately or specific to each demographic group. To identify whether or not air humidity would influence hospitalization rates for respiratory diseases, we obtained average annual rainfall data from municipalities in Climate-Data (Climate-Data, 2019).

Preliminarily, we tested for correlation among our explanatory variables to remove highly correlated variables (|r| ≥ 0.65) from our main analyses. For this reason, it was not possible to use the same units in all landscape metrics. The non-correlated explanatory variables (|r| ≤ 0.65) (Table 1) were included in the generalized linear models (GLM), which had as variables responses the rates of human admissions for respiratory diseases or for mental illness. After assessing the normality and homoscedasticity of the residuals from the most parameterized models in each analysis, we specified the Gaussian distribution family with the identity link function to perform the GLM’s. Using Akaike information criterion (AIC), in which each explanatory variable was associated with a model with a response variable (rate of hospitalization for mental illness or respiratory diseases), we considered the variable(s) of the best ranked models (Δ AIC ≤ 2) as the most likely determinants of influencing the hospitalization rates (Burnham and Anderson, 2002). We used the AICcmodavg package (Mazerolle, 2017) from the RStudio Program (v. 1.1.442) for analyses.

**Table 1:**
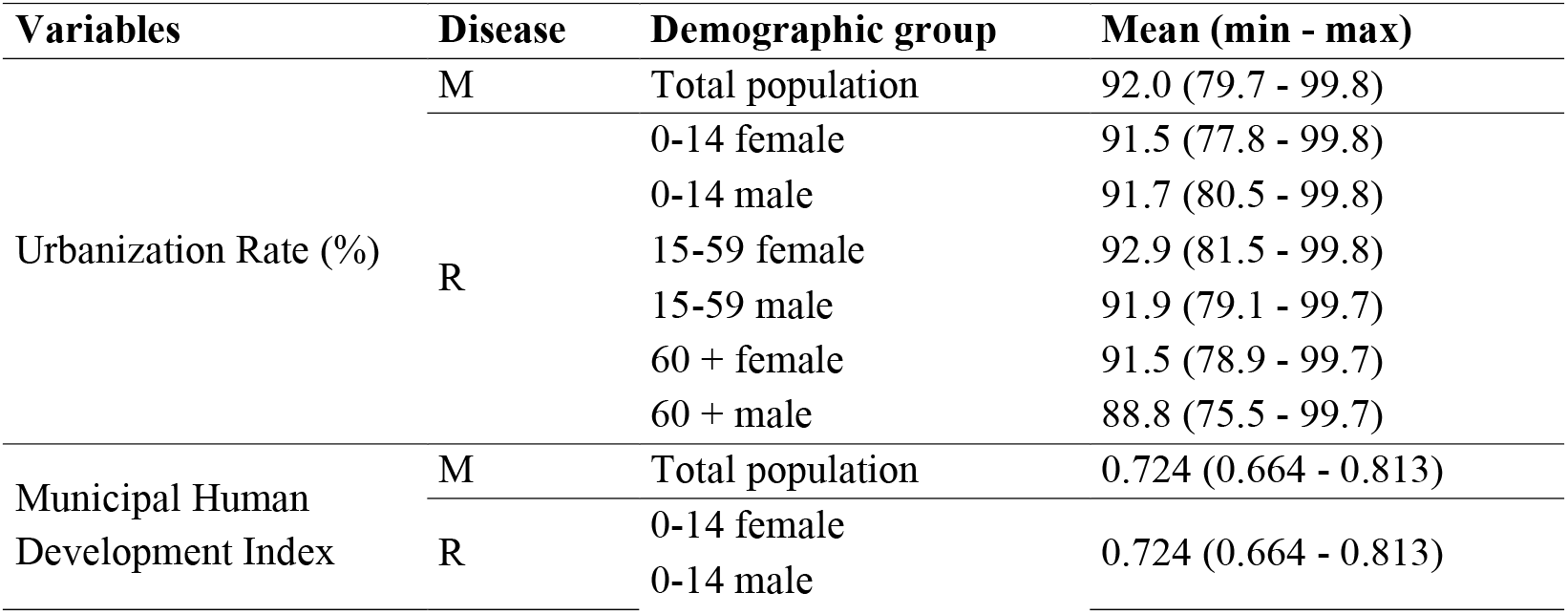
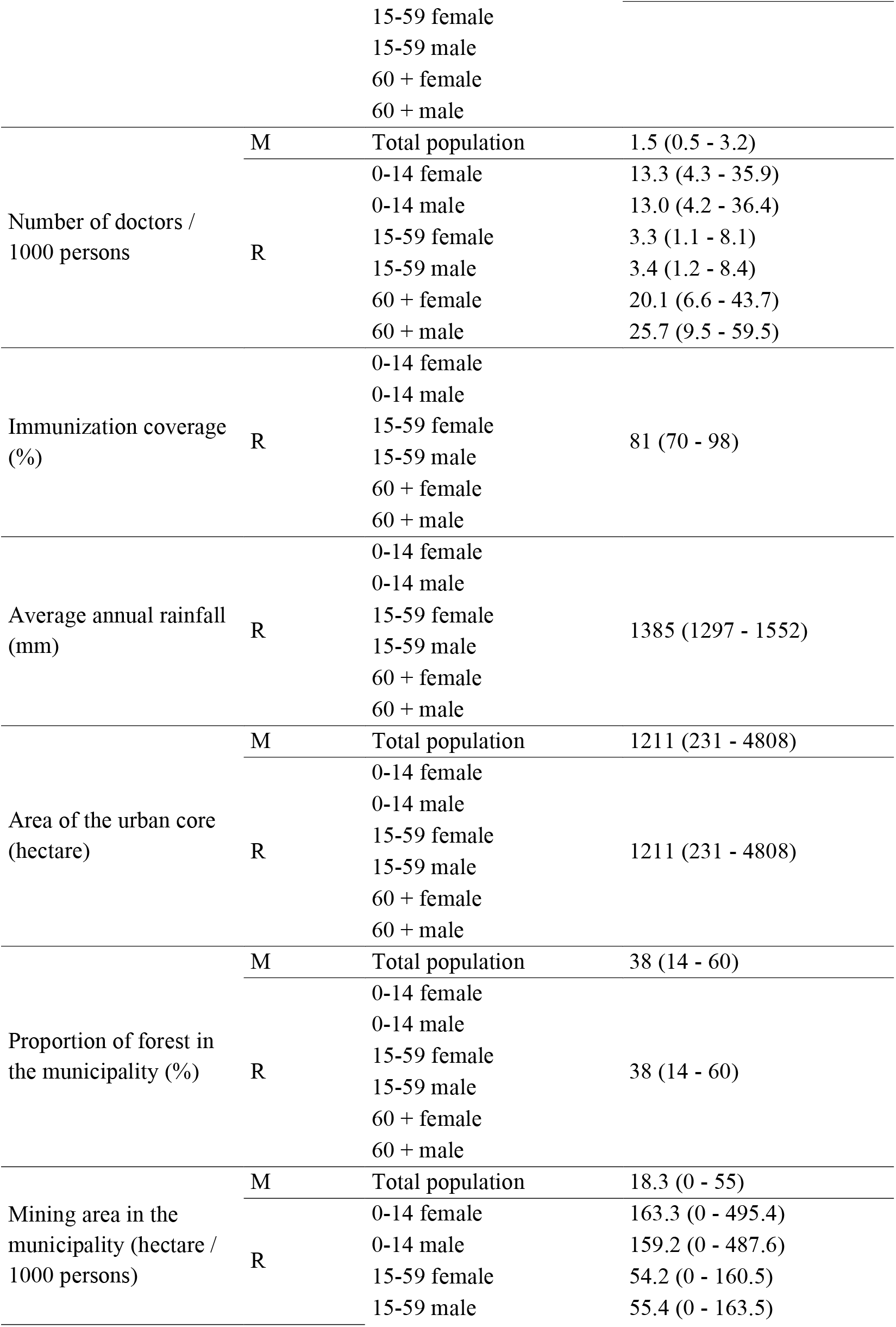
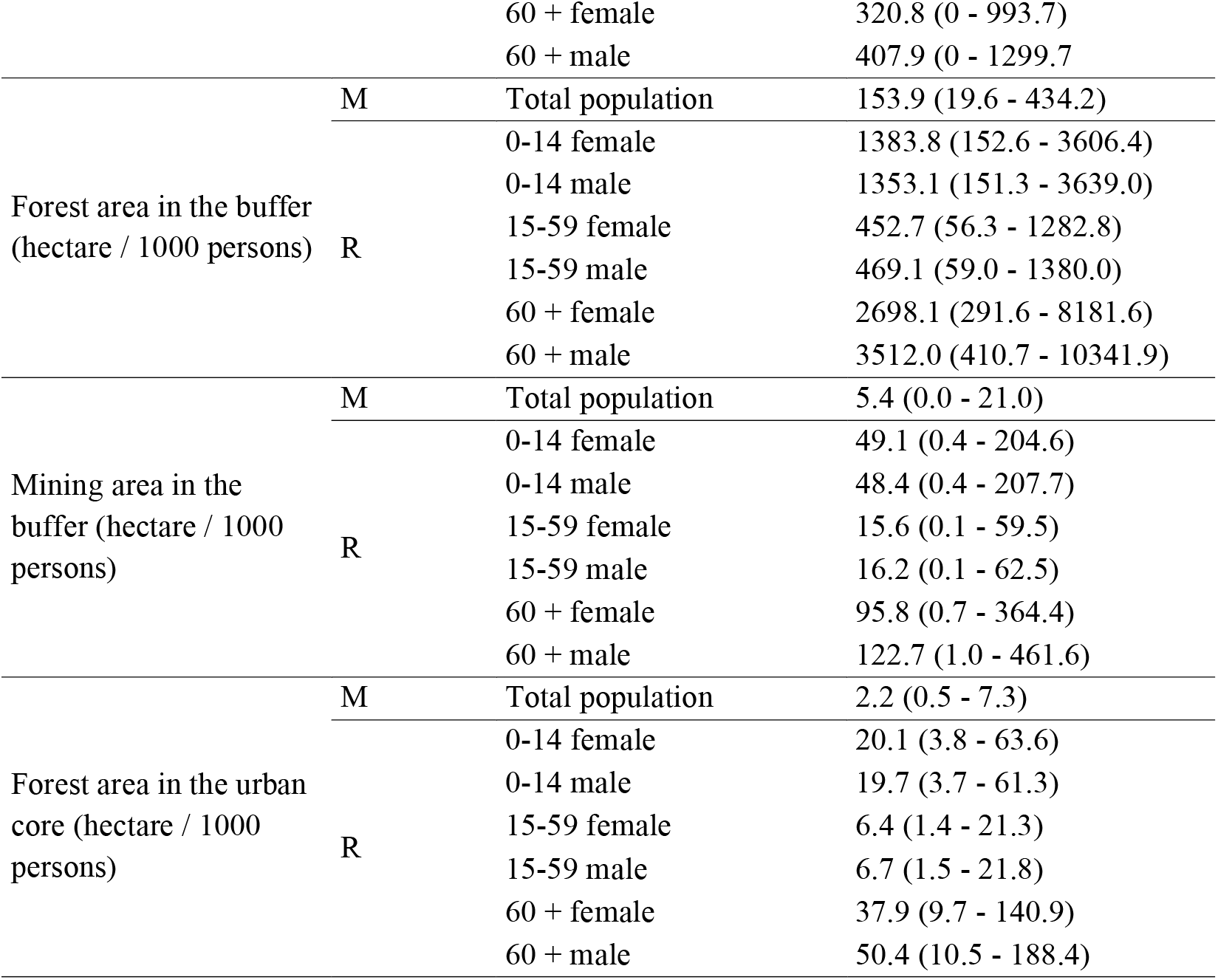
Socioeconomic and environmental variables used in the model selection analysis to assess the factors influencing hospitalization rates for respiratory diseases and mental illness in 22 municipalities in the Quadrilátero Ferrífero, state of Minas Gerais, Southeastern Brazil. M = Mental and behavioral disorders (i.e., mental illness); R = Respiratory diseases.

## Results

The number of hospitalizations for mental illness ranged from zero to 79 among municipalities, with an average of 16.8 (SD = 21.2) hospitalizations per municipality in 2010. Overall, there were 369 hospitalizations for mental illness in the 22 municipalities, an incidence of 31 hospitalizations / 100 thousand inhabitants, with an average of 11.4 (SD = 20.5) days of hospitalization per patient. Considering only the direct impacts, SUS had a cost of R$ 204,106.80 (R$ 340,494.21 in adjusted values) in the period. Among the nine variables analyzed, only the rate of urbanization influenced the rate of hospitalization for mental illness among the target municipalities. Municipalities with higher rates of urbanization had higher rates of hospitalizations for mental illness (Table 2 and Figure 2).

**Table 2:**
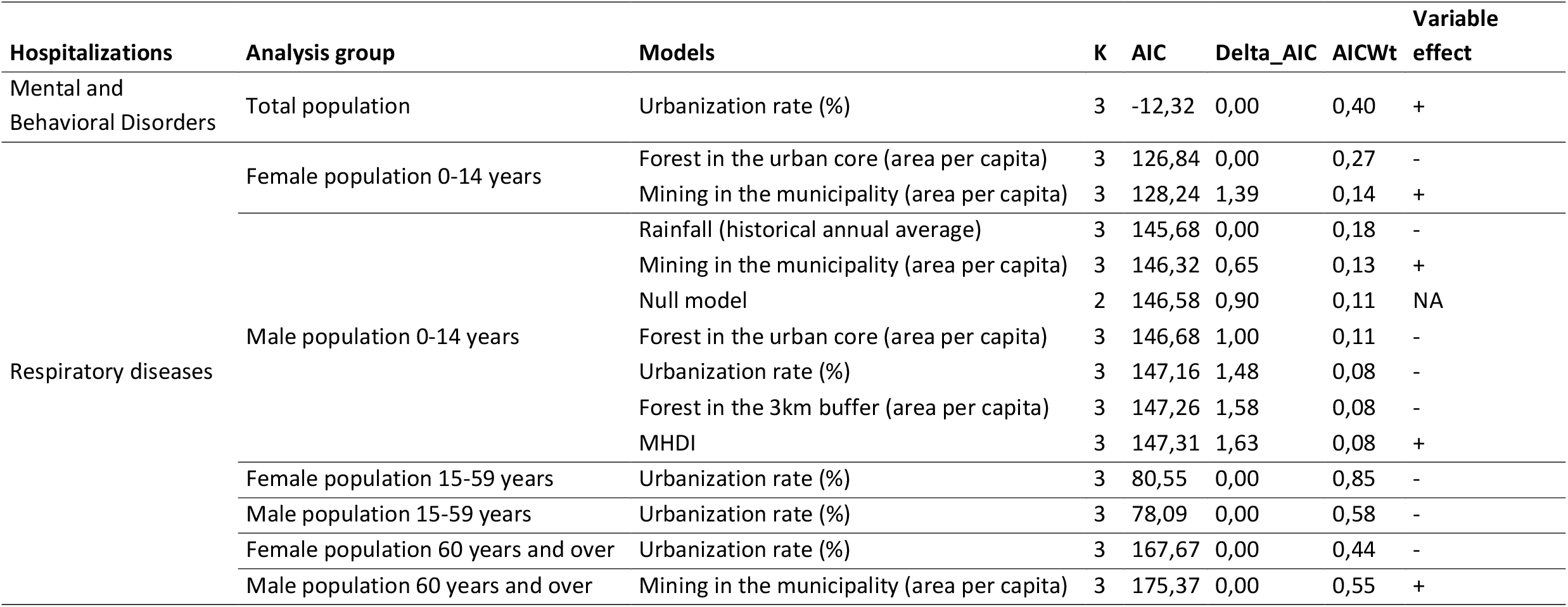
Model selection results containing the socioeconomic and environmental variables that influenced (Δ AIC ≤ 2) the hospitalization rates for respiratory diseases and for mental and behavioral disorders (i.e., mental illness) in 22 municipalities in the Quadrilátero Ferrífero, state of Minas Gerais, southeastern Brazil. + = positive effect of the explanatory variable on the response variable; - = negative effect of the explanatory variable on the response variable; NA = Not applicable; AICWt = model weight.

**Figure 2:**
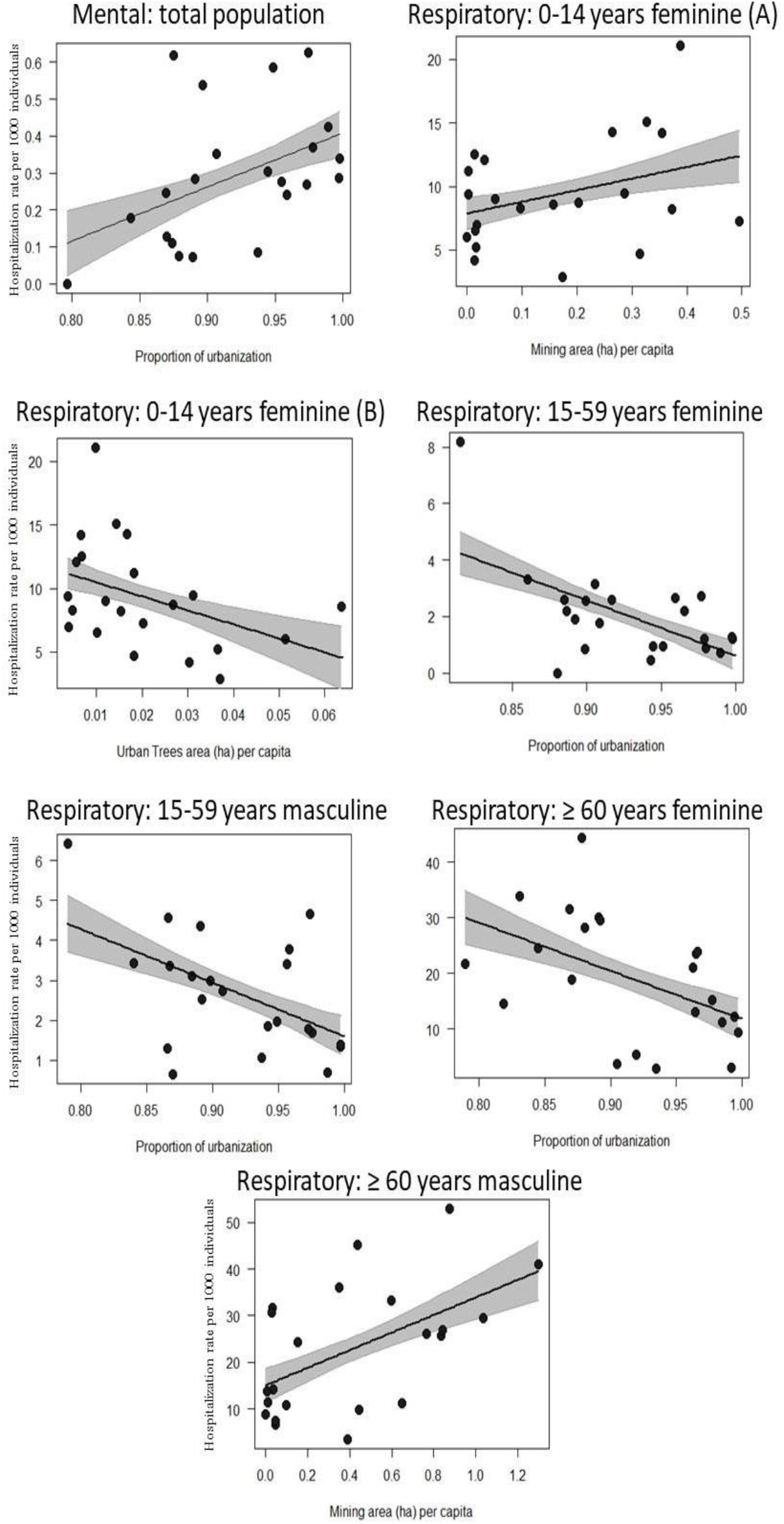
Estimates (± 95% CI) of the effect of explanatory variables (Δ AIC ≤2) on hospitalization rates for mental illness and respiratory diseases in the different demographic groups evaluated in 22 cities in the Quadrilátero Ferrífero, state of Minas Gerais, southeastern Brazil. A and B represent different variables that influenced the same socio-demographic group.

The number of hospitalizations for respiratory diseases ranged from 16 to 1,074 among the municipalities, with an average of 326 (SD = 296.9) hospitalizations per municipality in 2010. Overall, there were 7,176 hospitalizations in the 22 municipalities analyzed, at an incidence of 602 hospitalizations / 100 thousand inhabitants, with an average of 6.2 (SD = 1.5) days of hospitalization per patient. The Health System alone had a direct cost of R$ 7,090,041.57 (R$ 11,827,720.08 in adjusted amounts) in the period.

Considering the population from 0 to 14 years old, 1,422 female patients were hospitalized (1,033 hospitalizations / 100 thousand inhabitants), with an average of 3.9 (SD = 1.2) days of hospitalization and direct cost of R$ 991,851.22 (R$ 1,654,621.98 in adjusted amounts) for SUS, while 1,840 male patients were hospitalized (1,303 hospitalizations / 100 thousand inhabitants), with an average of 4.5 (SD = 1.2) days of hospitalization and direct cost of R$ 1,586,897.82 (R$ 2,647,288.17 in adjusted amounts) to SUS. The rate of hospitalization of the young (0-14 years) female population due to respiratory diseases was lower in municipalities with a larger area of forest per capita in urban centers and higher in municipalities with a larger area per capita of mining (Table 2 and Figure 2). However, this relationship was not strong for the male gender of this age group, since the null model was supported by the data (Table 2).

Considering the population aged 15 to 59 years, 713 female patients were hospitalized (175 hospitalizations / 100 thousand inhabitants), with an average of 7.0 (SD = 2.7) days of hospitalization and a direct cost of R$ 669,064.76 (R$ 1,116,144.47 in adjusted values) for SUS, while 951 male patients were hospitalized (243 hospitalizations / 100 thousand inhabitants), with an average of 8.0 (SD = 3.2) days of hospitalization and direct cost of R$ 1,239,686.59 (R$ 2,068,064.88 in adjusted values) for SUS. The rate of hospitalization for respiratory diseases in the adult population (15-59 years), both female and male, was significantly lower in municipalities with a higher rate of urbanization (Table 2 and Figure 2).

Considering the population aged 60 or over, 1,186 female patients were hospitalized (1,856 hospitalizations / 100 thousand inhabitants), with an average of 7.9 (SD = 6.0) days of hospitalization and a direct cost of R$ 1,336,478,16 (R$ 2,229,534.12 in adjusted values) for SUS, while 1,064 male patients were hospitalized (2,173 hospitalizations / 100 thousand inhabitants), with an average of 8.1 (SD = 3.4) days hospitalization and direct cost of R$ 1,266,065.73 (R$ 2,112,070.98 in adjusted values) for SUS. The rate of hospitalization for respiratory diseases in the elderly (60 years or more) female population was lower in municipalities with a higher rate of urbanization, while the rate of hospitalization for male respiratory diseases of the same age group was higher in cities with a larger area *per capita* of mining (Table 2 and Figure 2).

## Discussion

Our study demonstrates that urbanization is associated with an increase in the number of hospitalizations for mental illness, suggesting that life in cities may be a contributing factor for worsening mental health of the population. Although a large number of factors, such as economic, social, physiological, environmental, psychological, behavioral, genetic and even epigenetic issues can influence mental health (Galea et al., 2011; Meyer-Lindenberg, 2014), the role of nature deserves to be highlighted for providing psychological ecosystem services for human beings (Bratman et al., 2019).

The greater contact and diversity of experiences in natural environments are related to a set of benefits for people’s mental health, contributing to the reduction of the risk factors for mental and behavioral disorders and other diseases (Bratman et al., 2019; Frumkin et al., 2017). However, there is a tendency to reduce the quantity and quality of these experiences by people around the world (Bratman et al., 2019; Frumkin et al., 2017). This reduction is often related to life in urban environments, and the consequent reduction in opportunities and less active search for contact with nature by people (Cox et al., 2018), with a downward trend over human generations (Soga and Gaston, 2016).

Regions with higher rates of urbanization tend to have worse mental health indicators, presenting higher rates of depression when compared to rural populations, which have greater contact with nature (Cox et al., 2018). Studies have identified an association between moving to rural environments and reducing the risk of schizophrenia, with the authors associating this with epigenetic factors, which could be a possible explanation for the negative effect of urbanization on mental health (Galea et al., 2011). The increase of natural areas in urban environments can be a way to mitigate stress (Triguero-mas et al., 2017) and mood and anxiety disorders (Nutsford et al., 2013), contributing to the improvement of mental health’s indicators.

Contact with nature seems to be especially important for young people, highlighting the importance of investigating specific relationships for different genders and age groups of the population (Piccininni et al., 2018). Living in rural settings, as well as increasing the minimum time of exposure to green areas, reduced the risk of attention deficit and hyperactivity disorder in children in New Zealand (Donovan et al., 2019), while greater contact with nature was the factor responsible for reducing hospitalizations for mental disorders among female adolescents in Canada (Piccininni et al., 2018). Despite the increase in the coverage of the Psychosocial Care Centers (CAPS) having been identified as responsible for reductions in psychiatric hospitalizations in Brazil (Miliauskas et al., 2019), only half of the municipalities analyzed had any CAPS in 2010, so it is unlikely that this factor influence the relationship between greater urbanization and higher rates of hospitalization for mental illness that we found.

A study in England with five environmental variables found that the abundance of birds in the evening and the vegetation cover in the neighborhood contribute to the reduction of stress, depression, and anxiety (Cox et al., 2017). There are thresholds for the proportion of vegetation cover from which these green areas promote these benefits, being 20% for stress and depression and above 30% for anxiety (Cox et al., 2017). In our study, we did not identify the influence of urban forests on the population’s mental health. One possible explanation is that our municipalities had low vegetation cover, with an average of 7.34% (SD = 5.25) of forest cover in urban centers, values far below the thresholds described by Cox et al. (2017). Regarding the percentage of forested areas in the municipality, a study in the Netherlands (Helbich et al., 2018) indicated thresholds (79% of the municipality’s area) from which green spaces provide greater benefits for mental health. Again, this threshold falls well above the values evaluated in the present study (Table 1).

Although medical availability varied between municipalities (Table 1), it was not associated with a reduction in hospitalization rates. It is noteworthy that this data refers only to the number of doctors in the municipality, with no information on the working time that each doctor dedicates to serving the population, nor the location of clinics and hospitals (i.e. if they are distributed in the municipality or concentrated in certain regions), factors that may be important to determine preventive effects of medical assistance and, consequently, influence the hospitalization rates. The Municipal Human Development Index (MHDI) showed little variation between the municipalities evaluated (Table 1), indicating a similarity in average per capita income, education, and longevity in the region and, consequently, less influence of this variable on the health of the QF population.

Regarding the influence of urbanization on the population’s respiratory health, a positive association between urbanization and reduced rates of hospitalization for respiratory diseases in adults (15-59 years) of both genders and elderly women was observed. There is no consensus in the literature on the influence of urbanization on the incidence of respiratory diseases. Siddharthan et al. (2019) compared the urban and rural population in Uganda, identifying a higher prevalence of chronic obstructive pulmonary diseases in the rural population, due to the greater use of plant biomass in daily life (i.e. burning for food preparation), and a higher prevalence of asthma in the urban population (Siddharthan et al., 2019). A recent review (Sciaraffa et al., 2017) showed a correlation between urbanization and worse air quality indexes, with a higher frequency of asthma and respiratory infections, when comparing rural and urban populations in Italy. It should be noted that the municipalities evaluated in our study are small and medium-sized (the total population ranged from 3,513 to 202,942 people), which may be related to lower levels of air pollution resulting from the burning of fossil fuels in the road fleet.

Factors related to preventive health, such as ease of physical and logistical access to medicines, pharmacies, and hospitals, may explain the lower rates of hospitalization observed in municipalities where people live more in urban than rural environments. In a study conducted in the United States, there was a greater presence of chronic obstructive pulmonary diseases in the rural population compared to the urban population (Croft et al., 2018), which can be explained by the difference in access to health systems between populations, due to the greater reluctance of the rural population to seek preventive medical assistance, less availability of medical services in the rural area and greater difficulty for the rural population to access these services (Douthit et al., 2015). The greater density of pharmaceutical establishments, with the consequent higher availability of medicines, are related to a reduction in hospitalization rates for chronic diseases in Brazil through the Popular Pharmacy Program (PFP), as well as the city’s exposure time to this Federal Supply Program of free or low-cost medication (Almeida et al., 2019). The increase in the availability of medical assistance and prophylactic measures in rural areas, aiming at reducing the isolation of the population, may bring a consequent reduction in the rates of hospitalizations for respiratory diseases.

Several studies show the association of vaccination with a reduction in the incidence of respiratory diseases (Griffin et al., 2013; Silva et al., 2016). However, vaccination coverage showed little variation between municipalities (mean = 81.4%, SD = 8.9), so it was not feasible to assess the association between this variable and the rate of hospitalizations for respiratory diseases. Likewise, the municipalities presented values close to rainfall (mean = 1,385 mm, SD = 57), indicating a climatic similarity between the evaluated municipalities and, therefore, a lack of association between this variable and the rates of hospitalizations for respiratory diseases.

The reduction in hospitalization rates for respiratory diseases for the young population (female gender 0-14 years old), observed in our study in municipalities with greater tree vegetation coverage in urban centers, can be explained by the better air quality in municipalities with more vegetation in the urban area and / or factors related to the increase in population immunity due to contact with greater biodiversity of microorganisms. The fact that the null model was ranked together with other models for the young male population aged 0-14 years old, generates uncertainties about the factors that influence hospitalization rates for this group. Thus, further studies are needed to verify whether the same factors that determine hospitalization rates for the young female population are also determinant for the young male population.

Urban trees and shrubs are responsible for a set of ecosystem services, among which are the improvement of air quality by removing pollutants (O_3_, PM_10_, NO_2_, SO_2_, and CO) through the absorption of gases by leaf stomata and interception of particulate matter by leaves and branches (Nowak et al., 2006).. In a study carried out in the United States, the ecosystem service for removing five pollutants from the air by urban trees and shrubs was estimated at 711 thousand tons, valued at the time at U$ 3.8 billion annually (Nowak et al., 2006), demonstrating the importance of estimating and valuing this ecosystem service and allowing these externalities generated by different sectors of society to be internalized in the economy (Nowak et al., 2006).

Another mechanism that could explain the results found on respiratory health is related to the regulation and training of the immune system, dependent on contact with the biodiversity of microorganisms (Mills et al., 2019; Ruokolainen et al., 2015), especially in the early stages of life (Ruokolainen et al., 2015). Urbanization can reduce individual’s contact with microorganisms in the environment, leading to an increase in diseases such as asthma and allergies (Mills et al., 2019). The maintenance and increase of urban green areas, especially those with greater biodiversity, can contribute to the reduction in disease rates (Mills et al., 2019), consequently contributing to lower rates of hospitalization for respiratory diseases.

Urban forests could be transformed into parks or protected areas, and the financial resources for the creation and maintenance of these areas could come either from the secretariats / ministries of the environment, or from the secretariats / ministries of health, as it is an infrastructure of preventive health care. The creation of these urban parks must follow strategies that combat environmental injustices, ensuring that the entire population, especially those most vulnerable socioeconomically, can benefit from these ecosystem services, without, however, boosting real estate speculation and the phenomenon of gentrification (Wolch et al., 2014).

The impact of mining on the health of the young population (female gender 0-14 years old) is consistent with another study carried out in Itabira, a mining town in the QF, which related the increase in the attendance by acute air pollution of children and adolescents in emergency rooms with higher rates of inhalable particulate matter (PM10) from iron mining (Braga et al., 2007). Our study demonstrates that this relationship may be a pattern in municipalities that present open iron mining, evidencing the need to adopt mitigating actions to reduce particulate materials from these enterprises, aiming to reduce the impact on the health of children and adolescents by respiratory diseases. Impacts on the population’s respiratory health, related to mining activities, have also been identified in other regions of the world, with an increase in chronic bronchitis, emphysema and other chronic obstructive pulmonary diseases in the United States (Mabila and Almberg, 2018) and Sweden (Hedlund et al., 2004).

Air quality in mining cities is negatively affected by the emission of particulate material and aerosols from the activities of extraction, transportation, processing, and storage of iron ore and the waste and tailings generated (Csavina et al., 2012; Tavares et al., 2017). Particulate matter from open-pit mines has the potential to reach the entire urban area of the municipality, contributing directly and/or indirectly to air pollution and impacting the population’s health (Braga et al., 2007; Csavina et al., 2012; Tavares et al., 2017). Climatic factors can exacerbate these effects, allowing efflorescent deposits, with concentrations of metals and metalloids higher than the rest of the tailings, to be eroded from the tailing dams and carried by the wind over long distances, which may represent an additional risk to the nearby population’s health (Csavina et al., 2012). Open-pit mining, therefore, presents the greatest potential risk to human and environmental health among natural and anthropogenic sources of air particulate emissions (Csavina et al., 2012). It is essential to monitor and analyze, in a segmented manner, the concentrations of heavy metals in particulate material from the activities of the mines, considering phases of extraction, transportation, processing, and deposition of tailings independently, to determine the correct control measures.

In our study, as the male elderly population (60 years old or more), but not the female elderly, was influenced by the mining area of the municipality, it may indicates that this increase in hospitalizations is related to occupational activities that occurred during the mining operations that have historically occurred in the QF region. It is well-known that there is a predominance of male employees in mining activities (Mabila and Almberg, 2018), activities more susceptible to respiratory diseases (Hedlund et al., 2004; Mabila and Almberg, 2018). Other behavioral factors may also have influenced this result, such as, for example, the higher prevalence of smoking in the male population compared to the female population in Brazil (INCA, 2019).

## Conclusion

Given our data, we can conclude that mining activity is associated with a deterioration in health and, consequently, in the quality of life of the inhabitants of the municipalities of one of the largest mining provinces in the world. Our results indicate specific research paths to assess cause-effect relationships and demonstrate the importance of accounting for the ecosystem services, the depreciation of natural capital, and the negative impacts of landscape modifications in the evaluation of the cost-benefit of large enterprises. They also demonstrate the importance of population segmentation at different ages and genders and of assessing the landscape at different scales to determine the factors that influence people’s health. The method developed for this work can be replicated for other regions of Brazil and the world, having proved to be efficient in determining the socioeconomic environmental factors that contribute to collective health in countries and regions with a database geographically limited to municipalities.

The creation and maintenance of urban parks is a public health policy that prevents the occurrence of respiratory diseases in the young population and should be encouraged and seen as an investment in the quality of life and reduction of expenses with hospitalizations and loss of economic productivity. The quantification and valuation of these externalities are essential for deciding on whether or not to carry out a particular project. However, it is worth noting that economic valuation *per se* is not capable of internalizing all nature’s contributions, especially those related to human well-being. Perhaps, more important than the economic impact is the immaterial value of this ecosystem service, reflected in the value of people’s lives and well-being.

## Acknowledgments

MCF thanks FAPEMIG for the master’s degree scholarship in Brazil. RLM thanks CAPES for the grants. FHGR thanks CNPq for research schollarship. All authors are grateful to FAPEMIG, CAPES and CNPq for support. This research did not receive any specific grant from funding agencies in the public, commercial, or not-for-profit sectors.

